# C4 photosynthesis with Kranz anatomy evolved in the *Oryza coarctata* Roxb

**DOI:** 10.1101/2020.07.12.199232

**Authors:** Soni Chowrasia, Tapan Kumar Mondal

**Affiliations:** ICAR-National Research Institute for Plant Biotechnology, LBS Centre, Indian Agricultural Research Institute, New Delhi, 110012

## Abstract

The C4 cycle is a complex biochemical pathway that has been evolved in plants to deal with the adverse environmental conditions. Mostly C4 plants grow in arid, water-logged area or poor nutrient habitats. Wild species, *Oryza coarctata* (genome type KKLL; chromosome number (4x) =48, genome size 665 Mb) belongs to the genus of *Oryza* which thrives well under high saline as well as submerged conditions. Here, we report for the first time that *O. coarctata* is a C4 plant by observing the increased biomass growth, morphological features such as vein density, anatomical features including ultrastuctural characteristics as well as expression patterns of C4 related genes. Leaves of *O. coarctata* have higher vein density and possess Kranz anatomy. The ultrastructural observation showed chloroplast dimorphism i.e. presence of agranal chloroplasts in bundle sheath cells whereas, mesophyll cells contain granal chloroplasts. The cell walls of bundle sheath cells contain tangential suberin lamella. The transcript level of C4 specific genes such as *phosphoenolpyruvate carboxylase, pyruvate orthophosphate dikinase, NADP-dependent malic* enzyme and *malate dehydrogenase* was higher in leaves of *O. coarctata* compare to high yielding rice cultivar (IR-29). These anatomical, ultra structural as well as molecular changes in *O. coarctata* for C4 photosynthesis adaptation might be might be due to its survival in wide diverse condition from aquatic to saline submerged condition. Being in the genus of *Oryza*, this plant could be potential donor for production of C4 rice in future through conventional breeding, as successful cross with rice has already been reported.

## Introduction

Most of the plant species rely on RuBsiCO (Ribulose-1,5-bisphosphate carboxylase) enzyme for carboxylation of atmospheric carbon dioxide to produce phosphoglyceric acid (Calvin et al., 1949). The adverse environmental conditions allow the evolution of another biochemical mechanism to fix carbon dioxide by phosphoenolpyruvate carboxylase (PEPC) to produce oxaloacetic acid (Ehleringer et al., 1997). Beside this biochemical evolution, there are two major anatomical characteristics that had been evolved in C4 plants. They are i) presence of special anatomy of leaf i.e. Kranz (ring/ wreath-like) anatomy where bundle sheath cells surround vascular bundles (Dengler and Nelson, 1999) and ii) chloroplast dimorphism i.e. mesophyll chloroplasts contain well developed grana and bundle sheath chloroplasts are either agranal or possess unstacked grana (thylakoids) (Laetsch, 1974). The C4 plants evolved around 20 to 30 million years ago (Kellogg, 1999). So, far C4 photosynthesis had been found in 21 taxonomical groups i.e. 18 dicots (2,994 species) and 3 monocots (11,900 species) (Sage at al., 2001). However, it has been suggested that C4 plants are more like to be tolerant to different abiotic stresses (Bromham and Bennett, 2014).

*Oryza* genus comprises of 24 species out of which two species are edible and rests are grown as wild species (Sanchez et al., 2013). The wild plant species are excellent source of stress tolerance genes (Mondal et al., 2018). However, rice is a C3 plant therefore several attempts have been made to transfer C4 genes from maize into rice for increasing yield as well as better tolerance of stress (Ku et al., 1999; Takeuchi et al., 2000). *O. coarctata* is a wild halophyte found in the coastal regions. It had ability to tolerate high salt stress condition (Electrical conductivity-E.Ce 20-40dSm^−1^) and submergence condition for 12h (Sengupta and Mujumder, 2010; Chowrasia et al., 2019; 2018). Recently the genome sequence of this wild species of rice has been decoded in our laboratory (Mondal et al., 2018). Several salt stress responsive genes from this plant had been characterized (Sengupta et al., 2009; Kizhakkedath et al., 2015). Moreover, attempts were made for introgression of salt stress responsive genes from *O. coarctata* into rice cultivar (Jena, 1994; Islam et al., 2017) through conventional crossing method. Although, *O. coarctata* is an obligate halophytic grass but no attempt was made to study its mode of photosynthesis. Moreover, it is well-known that stress tolerant plants found in arid and saline habitats are mostly C4 (Bromham and Bennett, 2014). Therefore, present study deals with biomass growth, anatomical and ultrastructural features as well as expression pattern of C4 related genes in the leaves of *O. coarctata* to understand the mode of photosynthesis.

## Methods

### Plant materials

The plantlets of *O. coarctata* was collected from coastal region of Sunderban, West Bengal and grown in net house of NIPB (National Research Centre for Plant Biotechnology), New Delhi under control condition in soil under 16h daylight and 8h dark, an irradiance of 56 μmol m^−2^ s^−1^ temperatures of 30°C, and a relative humidity of 60-70 %. The rice seeds of IR-29 were sown in soil and maintained under same condition. All sampling was done at morning between 10-11 am from two months old seedling.

### Anatomical study of leaf

The young leaves of two month old plants were collected from both IR-29 and *O. coarctata* for anatomical studies. Fine sections of leaf were cut with the help of a sharp razor to observe under light microscope (Nikon Digital Microscopes Eclipse 80i) at magnification of 40X and 100X. Three biological replicates were taken for analysis of vein density.

For ultrastructural observation, the young leaves of *O. coarctata* were sectioned to 1 mm^2^ size and fixed in a fixative (2% glutaraldehyde and 2% paraformaldehyde in 0.1M phosphate buffer pH-7) for 24h (Giuliani et al., 2013). The leaves sections were then treated with 1% (w/v) osmium tetroxide followed by standard serial dehydration of acetone for 15 min each and finally embedded in Spurr’s epoxy resin. The ultra thin leaf blocks were made by employing microtome (Leica, Model: RM2235, Germany) and stained with 4% (w/v) uranyl acetate and 2% (w/v) lead citrate. For observation of ultrathin blocks, transmission electron microscope (JEOL, Model: JEM-2100F, Japan) equipped with a Mega View III Digital Camera and Image J Imaging System was employed.

### qReal Time-PCR (qRT-PCR) analysis of C_4_ specific genes

To study the expression patterns of C4 specific genes such as *carbonic anhydrase (CA), PEPC, PPDK (pyruvate orthophosphate dikinase*), *NADP-dependent malic* enzyme (*NADP-ME*) and *malate dehydrogenase* (*MDH*), were selected from previous reports (Supplementary Table 1).

The young leaves of *O. coarctata* were collected from two months old plants for RNA extracted by Trizol reagent as per the manufacturer’s protocol (Invitrogen, Carlsbad, USA). The integrity of RNA was checked on formaldehyde-agarose gel (2% w/v). Around 1μg RNA was taken for cDNA preparation by employing PrimeScript 1st strand cDNA Synthesis Kit (Clonetech Takara Bio, USA) (Chowrasia et al., 2019). Detail of primers has been given in Supplementary Table 1.

The relative expression level of C4 genes were studied by using qPCR (Light Cycler 480 II, Roche Molecular Diagnostics, USA) (Chowrasia et al., 2019). The expression level of C4 genes were normalized with *eukaryotic initiation factor 4-α* (*eIF4-α*) as internal control by using 2^−ΔΔCq^ (Cq=threshold cycle) formula (Schmittgen and Livak 2008). Three biological replicates were taken for expression level analysis.

### Statistical analysis

All the differences in mean values of observed characteristics in both rice and *O. coarctata* were calculated by Student *t*-test at significance of P = 0.05 and 0.001 (Chowrasia et al., 2019).

## Result and discussion

Plants evolved several physiological and biochemical features to encounters different stresses condition. One such evolution in plants is change of mode of photosynthesis i.e. origin of C4 and CAM (Crassulacean Acid Metabolism) (Ehleringer and Monson, 1993). C4 mode of photosynthetic plants are found to be grown in adverse environmental condition with compair to C3 mode of photosynthetic plants (Sage et al., 1999). The growth and development of C4 plants were less affected by these harsh environmental condition because of higher rate of photosynthesis, efficient carbon dioxide fixation, lower rate of photorespiration and transpiration to check water loss (Bromham and Bennett, 2014). Rice is a C3 plant (Covshoff et al., 2016) and being in the same genus, we selected a high yielding but highly salt stress sensitive cultivar IR-29 for comparative study. Further, biomass of *O. coarctata* was found to be more than IR-29 (Fig. 1 A) which might be due to higher rate of photosynthesis in *O. coarctata* compare to IR-29. *O. coarctata* is a perennial grass which predominantly propagates vegetatively because seeds are recalcitrant in nature (Jagtap et al., 2006). The leaves of this wild species of rice are thicker, waxy and leathery (Figure 1B) to prevent transpiration loss.

**Figure 1.**
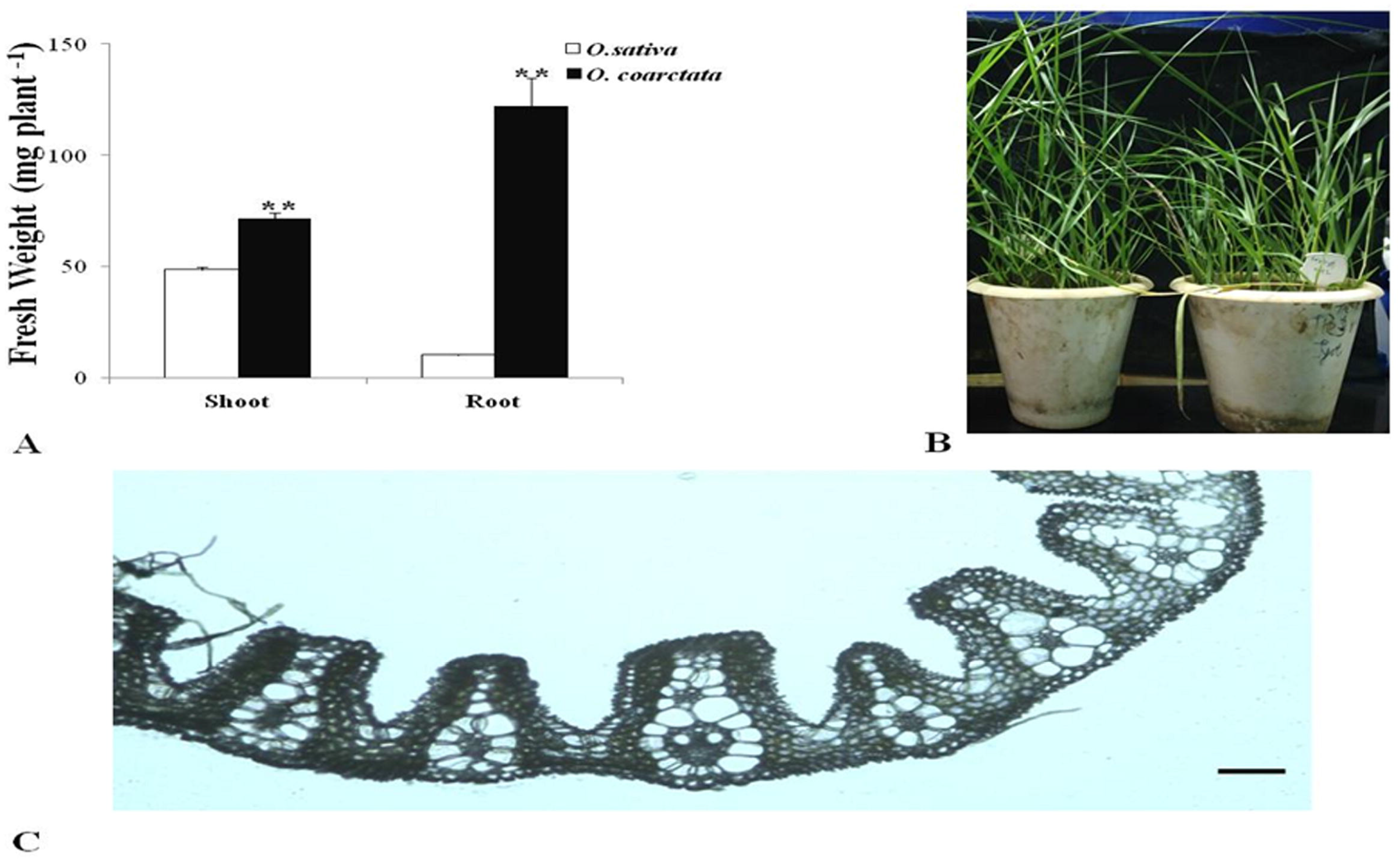
Morpho-anatomical study of *O. coarctata*. **(A)** Comparative observation of IR-29 (*O. saliva)* and *O. coarctata* biomass, **(B)** Vegetatively propagated seedling of *O. coarctata* and **(C)** Transverse section of young leaf of *O. coarctata* under light microscope (40X magnification). Scale bar=50μm. The means value of fresh weight (mg) were compared with t-test (** P ≤ 0.001).

Further, we studied the anatomical feathers of photosynthetically active two months old leaves of *O. coarctata* and IR-29. The transverse section of *O. coarctata* clearly shows Kranz anatomy i.e. wreath or ring like larger bundle sheath cells around the vascular bundle (Figure 1C). The Kranz anatomy is considered to be an exclusive characteristic of C4 plants (Lundgren et al., 2014). The vascular bundle of *O. coarctata* leaves is surrounded by the large bundle sheath cells which is a typical characteristics of C4 plant (Figure 1C) whereas, IR-29 being C3 plant showed anomalous non-Kranz anatomy (Weerasooriya et al., 2018). Like other C4 plants, bundle sheath cells were in turn surrounded by mesophyll cells. Each pair of vascular bundle is thus separated by two layers of bundle sheath and two layers of mesophyll cells (Figure 1C). The bundle sheath cells of *O. coarctata* are about 1.4 times larger than rice (Table 1). The leaf of *O. coarctata* was found to have many furrows and ridges, each ridge possess single vascular bundle. Therefore, the number of vein was also found to be higher in *O. coarctata* compare to IR-29 (Figure 2 A, B). For further detail study of veins, the leaves of both plant species were sectioned longitudinally. Both transverse (Figure 1C) and longitudinal (Figure 2C) sections of *O. coarctata* leaf blades showed large number of veins in compare to rice (Figure 2 B, D). In addition, the vein density of leaves was calculated and it was found that the distance between large transverse veins (TV), small longitudinal veins (SLV) and longitudinal veins (LLV) were 1.7, 1.56 and 1.86 times shorter in length in *O. coarctata* compared to rice (Table 1). This result depicts that large number of veins in leaves of *O. coarctata* might increases supply of water in leaves to maintain relative water content under saline condition. This increment of vein density in *O. coarctata* is one of the characteristic feathers of C4 plant (Kumar and Kellogg, 2019; Sage et al., 2004).

**Figure 2.**
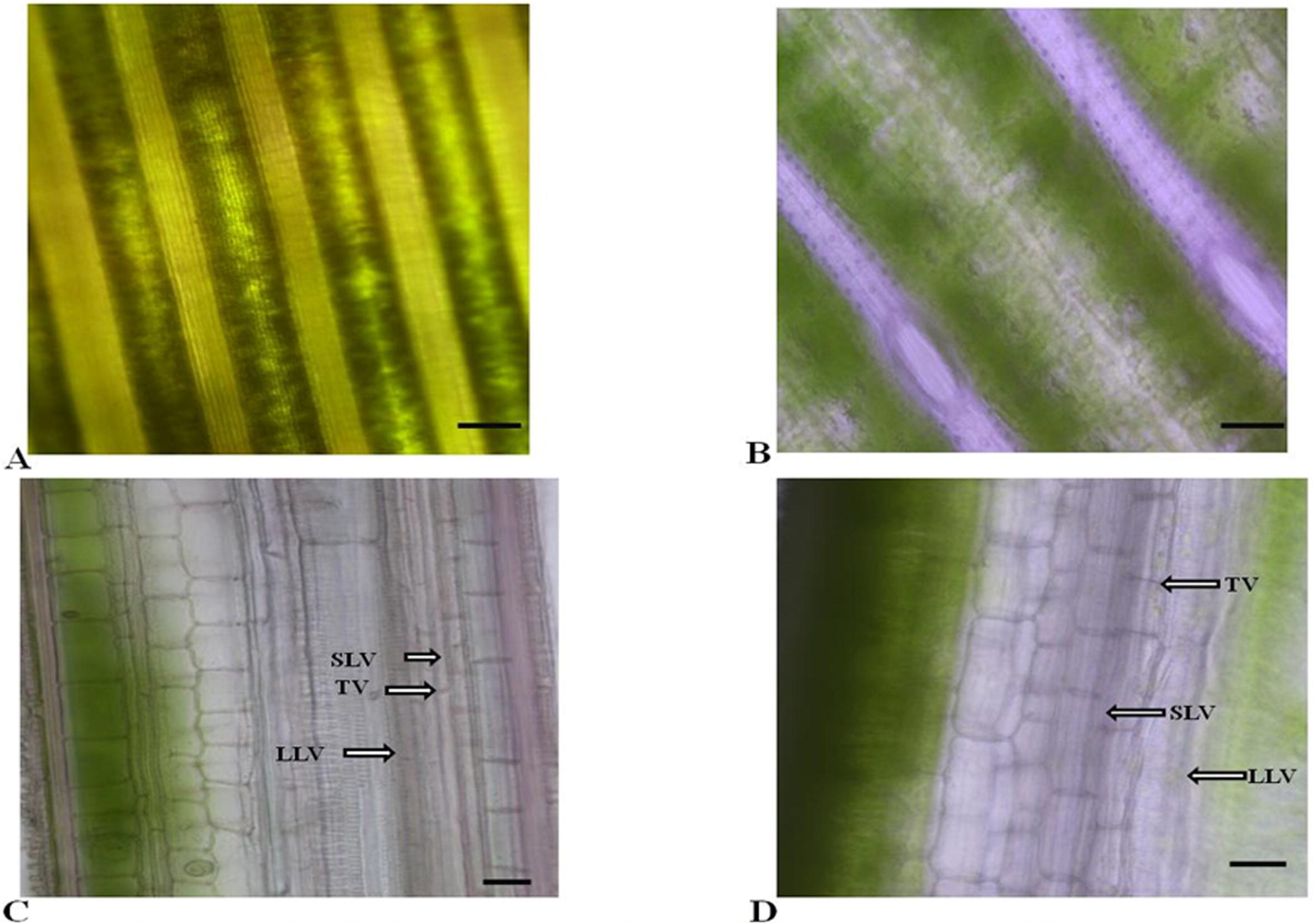
Images of leaf veins as observed under light microscope, **(A)** *O. coarctata* and **(B)** *O. sativa* (IR-29). **(C)** Longitudinal section of leaf blade of *O. coarctata* and **(D)** Rice. LLV-large longitudinal vein; SLV-small longitudinal vein and TV-transverse vein are indicated. Scale bar = 10μm.

**Table 1.** Detail study of leaf vein density. The value with same lower case are not significantly different in Student *t*-test at significance of *P* = 0.05 and 0.001. (*P*<0.001).

| Plant | Distance between large longitudinal veins ($\mu\text{m}$) | Distance between small longitudinal veins ($\mu\text{m}$) | Distance between transverse veins ($\mu\text{m}$) | Area of Bundle sheath ($\mu\text{m}^2$) |
| --- | --- | --- | --- | --- |
| *O. sativa* | 68.05<sup>a</sup> | 14.01<sup>a</sup> | 12.01<sup>a</sup> | 37.25<sup>a</sup> |
| *O. coarctata* | 40<sup>d</sup> | 8.94<sup>b</sup> | 6.60<sup>b</sup> | 52.97<sup>c</sup> |

The chloroplast dimorphism as well as peripheral localization of chloroplasts and mitrochondria are important characteristics of the C4 plant (Laetsch, 1974). To investigate chloroplast dimorphism and organelles arrangement in leaves *O. coarctata*, transmission electron microscopy was employed. It has been found that *O. coarctata* leaves exhibits typical arundinelloid type C4, where chloroplasts were present centrifugally in both mesophyll and bundle sheath cells. The mitochondrial number was higher as observed under TEM photograph in mesophyll cells compare to bundle sheath cells (Figure 3) which suggests slow rate of photorespiration in bundle sheath cells, an important adaptation of C4 plant (Kennedy et al., 1976). The outlines of bundle sheath and mesophyll cell wall were uneven and intercellar space was absent like other C4 plants (Dengler and Nelson, 1999) (Figure 3, 4). The mesophyll chloroplast possess granal stacking whereas, bundle sheath chloroplast lacks grana or contain rudimentary thylakoids (Figure 4, 5). The size of bundle sheath chloroplast was larger than mesophyll cells (Figure 5) which is one of the important criteria to depict that the C4 mode of photosynthesis of *O. coarctata*. The cell wall of bundle sheath cell was thicker than other mesophyll cells as shown in Figure 5 and suberin lamella was present on the cell wall of bundle sheath cell (Figure 6) which further indicates that *O. coarctata* possess C4 type of photosynthesis (Dengler and Nelson, 1999). Collectively, these ultrastructure characteristics of *O. coarctata* leaves revealed that it has adapted C4 mode of photosynthesis to deal with its adverse habitat (Muhaidat et al., 2011; Sage et al., 2004).

**Figure 3.**
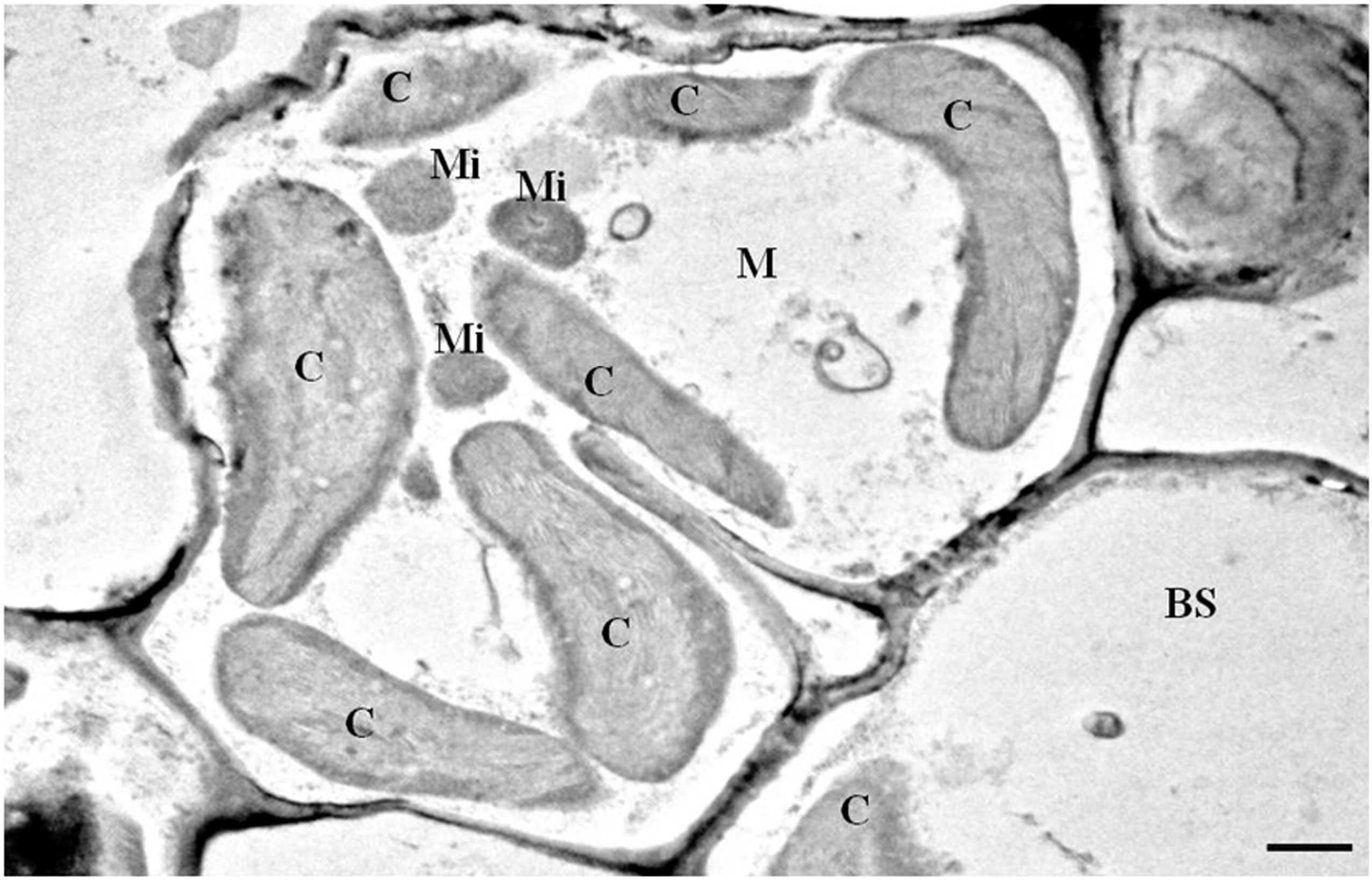
*O. coarctata* bundle sheath cell containing chloroplast and mitochondria. C-chloroplast, M-mesophyll, BS-bundle sheath and Mi-mitochondria. Scale bar = 500nm.

**Figure 4.**
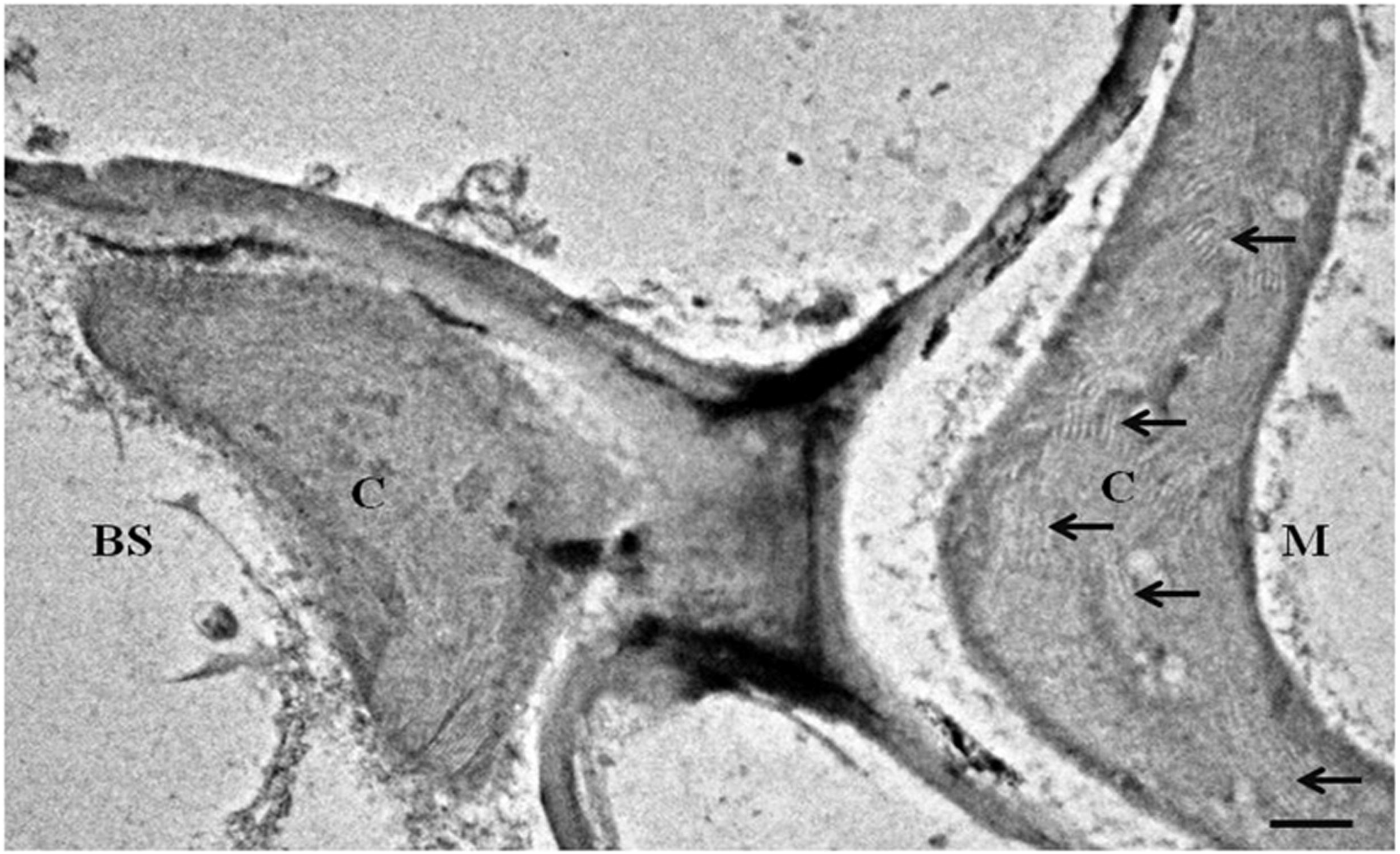
*O. coarctata* leaves exhibit chloroplast dimorphism i.e. mesophyll chloroplast exhibit extensive granal stacking (black arrow) and bundle sheath show no granal stacking under transmission electron microscope. Scale bar = 500nm.

**Figure 5.**
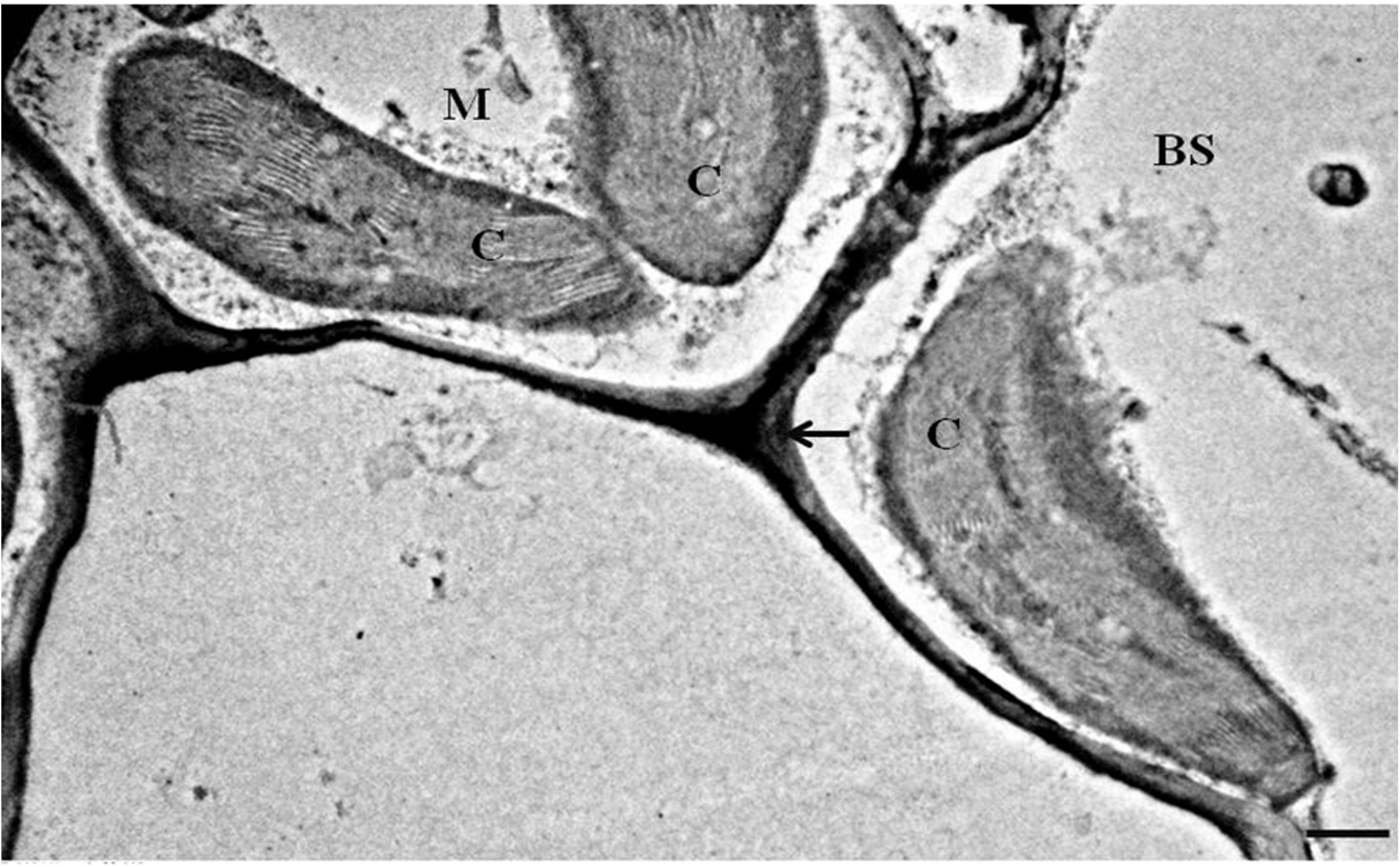
Ultrastructure of *O. coarctata* leaves illustrating centrifugal arrangement of the chloroplast in bundle sheath and mesophyll cells. The chloroplast of bundle sheath is larger than mesophyll chloroplast. The bundle sheath cell wall is thicker (black arrow). Scale bar = 500nm.

**Figure 6.**
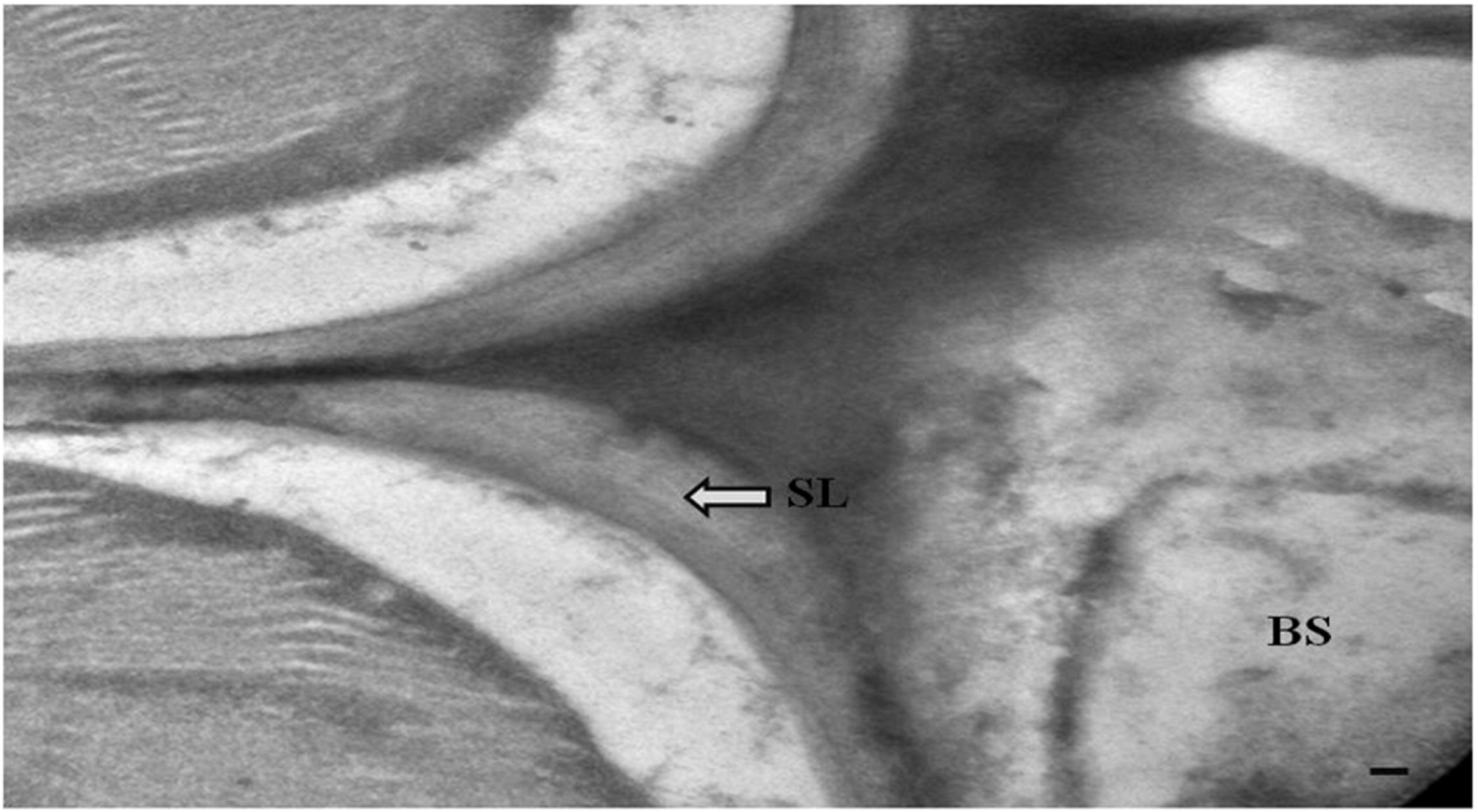
The bundle sheath cell wall showing suberin lamella (white arrow). Scale bar = 500nm. SL-suberin lamella.

In addition, we had also studied the relative expression level of C4 specific genes by RT-qPCR. Five C4 specific genes were selected based upon transcript abundance in C4 plants such as *Echinochloa glabrescens* (Covshoff et al., 2016) and maize (Xu et al., 2016) (Supplementary Table 1). The relative expression level of five C4 related genes i.e. *CA, PEPC, PPDK, NADP-ME* and *MDH* were compared in both leaves of *Oryza* species. The expression level of *CA* gene which is involved in hydration of carbon dioxide was found to be low in *O. coarctata* which corroborating with the findings of C4 plants (Xu et al., 2016). The transcript level of primary carboxylation gene of C4 cycle was significantly higher in *O. coarctata* compare to IR-29. Similarly, expression level of other important C4 related genes, *NADP-ME* and *MDH* were also found to be higher in *O. coarctata*. These two genes are involved in reduction of oxaloacetic acid into malate (Hatch and Slack, 1968). *PPDK* gene expression was also found to be high in *O. coarctata* compare to IR-29 rice variety, which involved in synthesis of phosphoenolpyruvate in chloroplast of mesophyll (Figure 7). Therefore, these above features depict that *O. coarctata* has C4 mode of photosynthesis.

**Figure 7.**
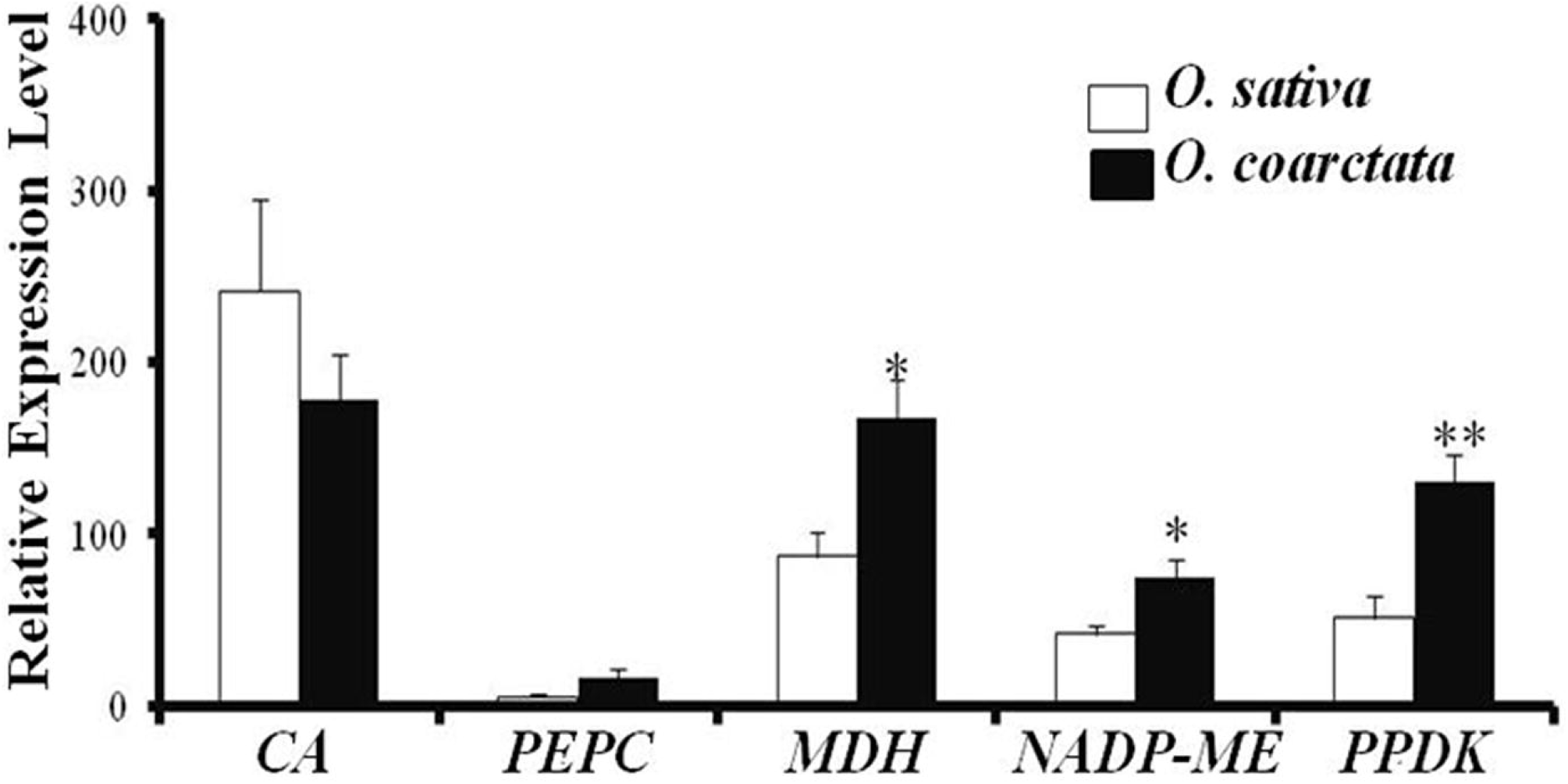
The relative expression levels of five C4 genes in rice and *O. coarctata as* normalized with *eukaryotic initiation factor 4-α*. The five C4 genes are *carbonic anhydrase (CA), phosphoenolpyruvate carboxylase (PEPC), malate dehydrogenase (MDH), NADP-dependent malic* enzyme *(NADP-ME)* and *pyruvate orthophosphate dikinase (PPDK)*. The means value of relative expression levels were with compared *t*-test (* P ≤ 0.05, ** P ≤ 0.001).

It was mainly found that C4 plants usually grown in warm temperate zone to tropical zone which consist of grassland, salt marshes (Sage et al., 1999). The high irradiance, low level of atmospheric carbon dioxide, high temperature, salinity, low nitrogen availability and drought promote C4 mode of photosynthesis (Osmond et al., 1982). Like other halophytic C4 plants, *O. coarctata* is shade loving and obligate halophyte where sodium is necessary for its growth (Sanchez et al., 2013). Sodium ions play an important role in pyruvate transportation through sodium:proton antiporter in plastid where it metabolized into carbon dioxide thus increasing its level for photosynthesis (Furumoto et al., 2011). Therefore, in order to scope such adverse condition (higher salt concentration E.Ce: 20-40 dSm^−1^) *O. coarctata* adapted efficient C4 mode of photosynthesis by expressing higher level of C4 specific genes and adapted special anatomical features to fix carbon dioxide.

Therefore, *O. coarctata* being halophyte has adapted C4 mode of photosynthesis mechanism among *Oryza* genus. Thus, introgression of C4 photosynthetic genes into rice by genetic engineering or by conventional breeding method will make C4 rice for higher yield.

## Acknowledgement

Authors are grateful to Mr. Sukhdev Nath, Swami Vivekananda Youth Cultural Society, Sagar, Sundarban, South 24 Parganas, West Bengal, India for supplying the *O. coarctata* plant and Prof Mohammad Anis, Department of Botany, Aligarh Muslim University, India and Dr. Kishore S. Rajput, Department of Botany, MS University Baroda, India for providing valuable suggestions for anatomical studies as well as Advanced Instrumentation Research Facility, Jawaharlal Nehru University, New Delhi, for Transmission electron microscopy study. SC acknowledges the financial support from Council of Scientific and Industrial Research, New Delhi. Authors are also grateful to Director, NIPB for providing the facility.

## Supplementary Table1

Gene specific primers of the selected C4 genes used in qRT-PCR.

## References

Bromham L. and Bennett T.H. (2014) Salt tolerance evolves more frequently in C4 grass lineages. J Evol. Biol. 27, 653–659. doi:org/10.1111/jeb.12320

Calvin M. (1949) The path of carbon in photosynthesis, VI. J Chem Educ 26, 639. https://doi.org/10.1021/ed026p639

Chowrasia S., Kaur H., Mujib A. and Mondal T.K. (2019) Evaluation of *Oryza coarctata* candidate reference genes under different abiotic stresses. Biol. Plant. 63, 496–503. doi: 10.32615/bp.2019.054

Chowrasia S., Rawal H.C., Mazumder A., Gaikwad K., Sharma T.R., Singh N.K. and Mondal T.K. (2018) *Oryza coarctata* Roxb. In The Wild Oryza Genomes (pp. 87–104). Springer, Cham.

Covshoff S., Szecowka M., Hughes T.E., Smith-Unna R., Kelly S., Bailey K.J., Sage T.L., Pachebat J.A., Leegood R. and Hibberd J.M. (2016) C4 photosynthesis in the rice paddy: insights from the noxious weed *Echinochloa glabrescens*. Plant Physiol. 170, 57–73. doi: 10.1104/pp.15.00889.

Dengler N.G. and Nelson T. (1999) Leaf structure and development in C4 plants. C. 4, 133–172.

Ehleringer J.R., Cerling T.E. and Helliker B.R. (1997) C4 photosynthesis, atmospheric CO2, and climate. Oecologia. 112, 285–299. doi: 10.1007/s004420050311

Furumoto T., Yamaguchi T., Ohshima-Ichie Y., Nakamura M., Tsuchida-Iwata Y., Shimamura M., Ohnishi J., Hata S., Gowik U., Westhoff P. and Bräutigam A. (2011) A plastidial sodium-dependent pyruvate transporter. Nature. 476, 472–475. doi: 10.1038/nature10250

Giuliani R., Koteyeva N., Voznesenskaya E., Evans M.A., Cousins A.B. and Edwards G.E. (2013) Coordination of leaf photosynthesis, transpiration, and structural traits in rice and wild relatives (genus *Oryza*). Plant Physiol. 162, 1632–1651. doi:https://doi.org/10.1104/pp.113.217497

Hatch M.D. and Slack C.R. (1968) A new enzyme for the interconversion of pyruvate and phosphopyruvate and its role in the C4 dicarboxylic acid pathway of photosynthesis. Biochem. J. 106, 141–146. doi: 10.1042/bj1060141

Islam T., Biswas S., Mita U.H., Sarker R.H., Rahman M.S., Ali M.A., Aziz K.S. and Seraj Z.I. (2017) Characterization of Progenies from Intergeneric Hybridization Between *Oryza sativa* L. and *Porteresia coarctata* (Roxb.) Tateoka. Plant Tissue Culture Biotech. 27, 63–76. doi.org/10.3329/ptcb.v27i1.35013

J.R. and Monson R.K. (1993) Evolutionary and ecological aspects of photosynthetic pathway variation. Ann. Rev. Eco. System. 24, 411–439. doi: 10.1146/annurev.es.24.110193.002211

Jagtap T.G., Bhosale S. and Charulata S. (2006) Characterization of *Porteresia coarctata* beds along the Goa coast, India. Aqua.Bot. 84, 37–44. doi:10.1016/j.aquabot.2005.07.010

Jena K.K. (1994) Production of intergeneric hybrid between *Orzya sativa* L. and *Porteresia coarctata* T. Current Sci. 67, 744–746.

Kellogg E.A. (1999) Phylogenetic aspects of the evolution of C4 photosynthesis. In C4 Plant Biology (Sage, R. F. and Monson, R. K., eds.), San Diego: Aca. Press. 12, 411–444.

Kennedy R.A. (1976) Photorespiration in C3 and C4 plant tissue cultures: significance of Kranz anatomy to low photorespiration in C4 plants. Plant Physiol. 58, 573–575.

Kizhakkedath P., Jegadeeson V., Venkataraman G. and Parida A. (2015) A vacuolar antiporter is differentially regulated in leaves and roots of the halophytic wild rice *Porteresia coarctata* (Roxb.) Tateoka. Mol. Biol. Rep. 42, 1091–1105. doi: 10.1007/s11033-014-3848-4

Ku M.S., Agarie S., Nomura M., Fukayama H., Tsuchida H., Ono K., Hirose S., Toki S., Miyao M. and Matsuoka M. (1999) High-level expression of maize phosphoenolpyruvate carboxylase in transgenic rice plants. Nature Biotech. 17, 76–80. doi.org/10.1038/5256

Kumar D. and Kellogg E.A. (2019) Getting closer: vein density in C4 leaves. New Phyto. 221, 1260–1267. doi.org/10.1111/nph.15491

Laetsch W.M. (1974) The C4 syndrome: a structural analysis. Ann. Rev. Plant Physiol. 25, 27–52. doi.org/10.1146/annurev.pp.25.060174.000331

Lundgren M.R., Osborne C.P. and Christin P.A. (2014) Deconstructing Kranz anatomy to understand C4 evolution. J. Exp. Bot. 65, 3357–3369. doi.org/10.1093/jxb/eru186

Mondal T.K., Rawal H.C., Chowrasia S., Varshney D., Panda A.K., Mazumdar A., Kaur H., Gaikwad K., Sharma T.R. and Singh N.K. (2018) Draft genome sequence of first monocot-halophytic species *Oryza coarctata* reveals stress-specific genes. Sci. Rep. 8, 1–3. doi: 10.1038/s41598-018-31518-y

Muhaidat R., Sage T.L., Frohlich M.W., Dengler N.G. and Sage R.F. (2011) Characterization of C3–C4 intermediate species in the genus *Heliotropium* L.(Boraginaceae): anatomy, ultrastructure and enzyme activity. Plant Cell Env. 34, 1723–1736. doi: 10.1111/j.1365-3040.2011.02367.x

Osmond C.B., Winter K. and Ziegler H. (1982) Functional significance of different pathways of CO 2 fixation in photosynthesis. In Physiological plant ecology II (pp. 479-547). Springer, Berlin, Heidelberg. doi.org/10.1007/978-3-642-68150-916

Sage R.F. (2001) Environmental and evolutionary preconditions for the origin and diversification of the C4 photosynthetic syndrome. Plant Biol. 3, 202–213. doi.org/10.1055/s-2001-15206

Sage R.F. (2004) The evolution of C4 photosynthesis. New Phyto. 161, 341–370.

Sage, R.F., Wedin, D.A. and Li, M., (1999) The biogeography of C4 photosynthesis: patterns and controlling factors. C4 Plant Biol, 1, 313–373.

Sanchez P.L., Wing R.A. and Brar D.S. (2013) The wild relative of rice: genomes and genomics. In Genetics and genomics of rice (pp. 9–25). Springer, New York, NY. doi.org/10.1007/978-1-4614-7903-1_2

Schmittgen, T.D. and Livak, K.J. (2008) Analyzing real-time PCR data by the comparative C_T_ method. Nature Protoc. 3, 1101–1108. doi.org/10.1038/nprot.2008.73

Sengupta S. and Majumder A.L. (2009) Insight into the salt tolerance factors of a wild halophytic rice, *Porteresia coarctata:* a physiological and proteomic approach. Planta. 229, 911–928. doi: 10.1007/s00425-008-0878-y

Sengupta S. and Majumder A.L. (2010) *Porteresia coarctata* (Roxb.) Tateoka, a wild rice: a potential model for studying salt stress biology in rice. Plant Cell Environ. 33, 526–542. doi: 10.1111/j.1365-3040.2009.02054.x.

Takeuchi Y., Akagi H., Kamasawa N., Osumi M. and Honda H. (2000) Aberrant chloroplasts in transgenic rice plants expressing a high level of maize NADP-dependent malic enzyme. Planta. 211, 265–274. doi.org/10.1007/s004250000282

Ueno O. and Sentoku N. (2006) Comparison of leaf structure and photosynthetic characteristics of C3 and C4 *Alloteropsis semialata* subspecies. Plant Cell Envir. 29, 257–268. doi: 10.1111/j.1365-3040.2005.01418.x

Weerasooriya H.A., Jayasekera A. and Caldera I. (2018) Tendencies towards a C4 Leaf: Quantitative Studies on Leaf Anatomy of Selected C3 and C4 Grasses. J Rice Res. 6, 2–9. doi:10.4172/2375-4338.1000188

Xu J., Bräutigam A., Weber A.P. and Zhu X.G. (2016) Systems analysis of cis-regulatory motifs in C4 photosynthesis genes using maize and rice leaf transcriptomic data during a process of deetiolation. J. Exper. Bot. 67, 5105–5117. doi.org/10.1093/jxb/erw275

