## Supplementary Table 1 for "C4 photosynthesis with Kranz anatomy evolved in the *Oryza coarctata* Roxb"

**Supplementary Table1**. Gene specific primers of the selected C4 genes used in qRT-PCR.

| **Gene name** | **Gene id** | **Forward primer sequence** | **Reverse primer sequence** | **Tm (melting temperature)** | **Product size (base pair)** | **References** |
| --- | --- | --- | --- | --- | --- | --- |
| Carbonic anhydrase | LOC_Os01g45274 | GCCAAGTTCAAGACCGAGTTCTATGACA | AGCGTGCTTGATCTTGCAGTAAGCTG | 61 | 204 | Covshoff et al., 2016. |
| Phosphoenolpyruvate carboxylase | LOC_Os01g11054 | GGAAGAAGATTTCTCCAGGAGAACCTTAC | CAAGAACTGCTCGACATTGGTGTAAGTC | 58 | 155 | Xu et al., 2016. |
| Malate dehydrogenase | LOC_Os08g44810 | GGAGACCAGTGAAAGAAGTCATTAAAGACAC | CATACTGAACACGATGTCCTCTGCTATG | 58 | 248 | Covshoff et al., 2016. |
| NADP-dependent malic enzyme | LOC_Os01g09320 | CTGATACAGTTTGAGGACTTCGCCAATC | GAGCAATGAGTTCTGCAATACCAGTTCC | 60 | 223 | Covshoff et al., 2016. |
| Pyruvate orthophosphate dikinase | LOC_Os05g33570 | GTGTAAATGATGCGTCCAAGATTGTAG | GTATCAGCATTTGCCATAACCTTGAG | 57 | 202 | Xu et al., 2016. |
| Eukaryotic initiation factor 4-α | MK123515 | CAGCAACTTGACTATGGATTGGTGGA | CATCCAGCACAAACATCTTAATGTGGTC | 60 | 270 | Chowrasia et al., 2019. |
